# Sustained upregulation of widespread hippocampal-neocortical coupling following memory encoding

**DOI:** 10.1101/2022.01.07.475348

**Authors:** Line Folvik, Markus H. Sneve, Hedda T. Ness, Didac Vidal-Piñeiro, Liisa Raud, Oliver M. Geier, Kristine B. Walhovd, Anders M. Fjell

**Affiliations:** Center for Lifespan Changes in Brain and Cognition, Department of Psychology, University of Oslo, Oslo 0373, Norway; Department of Diagnostic Physics, Oslo University Hospital, Oslo 0424, Norway; Department of Radiology and Nuclear Medicine, Oslo University Hospital, 0372 Oslo, Norway

**Author notes:** **Corresponding author:** Line Folvik, Department of Psychology, University of Oslo, Oslo 0373, Norway. **Email:**. **Author Contributions:** L.F., H.T.N, M.H.S, D.V-P, O.G., K.B.W, and A.M.F designed research; L.F., H.T.N, M.H.S, D.V-P, L.R, and A.M.F. performed research; M.H.S., O.M.G, K.B.W, and A.M.F. contributed new reagents/analytic tools; L.F., H.T.N, M.H.S, D.V-P, L.R, and A.M.F. analyzed data; L.F., H.T.N, M.H.S, D.V-P, L.R., K.B.W, and A.M.F wrote the paper. **Competing Interest Statement:** The authors declare no conflict of interest.

**Keywords:** hippocampus, reactivation, systems consolidation, encoding, fMRI

## Abstract

Systems consolidation of new experiences into lasting episodic memories involves interactions between hippocampus and the neocortex. Evidence of this process is seen already during early awake post-encoding rest periods. Functional MRI (fMRI) studies have demonstrated increased hippocampal coupling with task-relevant perceptual regions and reactivation of stimulus-specific encoding patterns following intensive encoding tasks. Here we investigate the spatial and temporal characteristics of these hippocampally anchored post-encoding neocortical modulations. Eighty-nine adults participated in an experiment consisting of interleaved memory task- and resting-state periods. As expected, we observed increased post-encoding functional connectivity between hippocampus and individually localized neocortical regions responsive to stimulus categories encountered during memory encoding. Post-encoding modulations were however not restricted to stimulus-selective cortex, but manifested as a nearly system-wide upregulation in hippocampal coupling with all major functional networks. The spatial configuration of these extensive modulations resembled hippocampal-neocortical interaction patterns estimated from active encoding operations, suggesting hippocampal post-encoding involvement by far exceeds reactivation of perceptual aspects. This reinstatement of encoding patterns during immediate post-encoding rest was not observed in resting-state scans collected 12 hours later, nor in control analyses estimating post-encoding neocortical modulations in functional connectivity using other candidate seed regions. The broad similarity in hippocampal functional coupling between online memory encoding and offline post-encoding rest suggests reactivation in humans may involve a spectrum of cognitive processes engaged during experience of an event.

**Significance statement:** Stabilization of newly acquired information into lasting memories occurs through systems consolidation – a process which gradually spreads the locus of memory traces from hippocampus to more distributed neocortical representations. One of the earliest signs of consolidation is the upregulation of hippocampal-neocortical interactions during periods of awake rest following an active encoding task. We here show that these modulations involve much larger parts of the brain than previously reported in humans. Comparing changes in hippocampal coupling during post-encoding rest with those observed under active encoding, we find evidence for encoding-like hippocampal reinstatement throughout cortex during task-free periods. This suggests early systems consolidation of an experience involves reactivating not only core sensory details but multiple additional aspects of the encoding event.

## Introduction

Memory systems consolidation refers to the transformation of experiences into longer-lasting episodic memories via hippocampal-neocortical interactions (1, 2). Animal research has shown that such stabilization of memory traces results from spontaneous reactivations of hippocampal-neocortical connectivity patterns, which can occur both during deep sleep (3) and awake periods of rest (4). Due to its spontaneous nature, it is difficult to achieve adequate experimental control of systems consolidation resulting from hippocampal replay. In recent years, however, task-free functional magnetic resonance imaging (fMRI) and magnetoencephalography (MEG) have been used successfully to investigate reactivation-related processes in awake humans non-invasively and with high spatial precision (5–7). Several fMRI studies have found experience-dependent alterations in resting-state functional connectivity (rsFC) between hippocampus and category-sensitive cortices after an encoding task compared to a pre-encoding baseline measure (8–11). While most rsFC studies have focused on hippocampal interactions with pre-defined, task-relevant perceptual regions, investigations in non-human primates (12), and recently in humans using MEG (13), have shown that hippocampal states associated with reactivation and memory consolidation coincide with activity modulations in large parts of the neocortex, also beyond sensorimotor and perceptual cortices. Currently, however, we do not know whether these extensive neocortical modulations are part of a system-wide upregulation of hippocampal functional connectivity during post-encoding consolidation. Alternatively, hippocampal functional modulations could be limited to category-selective cortex while engagement of non-sensory networks could occur via alternative pathways, e.g., mediated by the thalamus (14). Critically, hippocampal functional connectivity modulations during memory-relevant task-states involve large portions of the neocortex (15–21). If these broader hippocampal networks are coordinated similarly also during post-encoding rest, this would suggest state continuation into periods without systematic exposure to external stimuli – akin to system-wide maintenance or reactivation of memory-relevant interactions between brain regions.

In the present fMRI study, we characterized the systems-level changes in hippocampal interactions occurring in humans following an intense learning session. Younger and older adults (N = 89; Table 1) completed an intentional associative memory encoding task involving stimulus categories for which there exist established functional localizers. Reactivation-related changes in hippocampal-neocortical rsFC were estimated from resting-state periods taking place immediately before and after the encoding task as well as after a delay of 12 hours (Figure 1). We ran additional localizer-sequences to individually map object-, face-, and place-sensitive regions. Our first aim was to replicate previous reports of increased post-encoding rsFC between hippocampus and these category-sensitive perceptual regions. Importantly, we also tested for reactivation effects outside stimulus-selective cortex; both exploratory at the whole-brain parcel-level and at the level of established “canonical” resting-state networks. Post-encoding modulations in hippocampal rsFC were compared with functional connectivity patterns estimated from encoding and retrieval task-periods. Strong resemblance with encoding patterns would support an interpretation of ongoing reactivation (7). Higher similarity with retrieval patterns, on the other hand, could indicate rehearsal-like operations and would potentially invalidate an interpretation of post-encoding rsFC modulations as reflective of spontaneous consolidation processes in the awake state. Finally, the current sample consisted of younger and older adults. With few exceptions (22), the effect of age on awake systems consolidation processes remains unexplored. Age is among the strongest individual predictors of episodic memory function, and the memory decline observed in normal aging is related to changes in hippocampal structure and function (23). Therefore, throughout the analyses, we systematically tested whether awake hippocampal reactivation-like patterns were affected by participant age.

**Figure 1.**
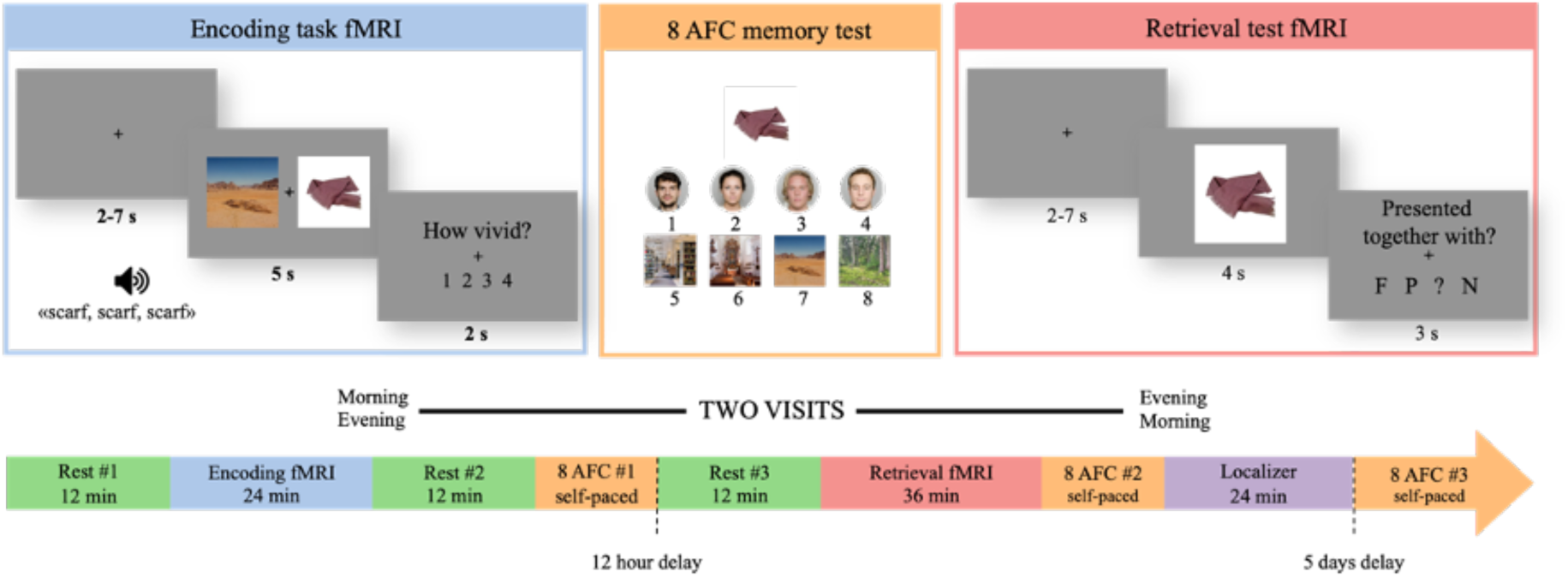
Experiment structure. Participants completed a multimodal associative encoding task, visualizing interactions between items and face/place associates. Memory for the encoded associations was tested via offline forced-choice retrieval after 15 minutes, 12 hours and 5 days, and during an in-scanner cued retrieval session taking place ∼12hours post-encoding. Here, a mix of previously encoded and novel items were presented and the participants indicated memory of the associate via F(ace) or P(place) responses, or alternatively item recognition without source information (“?”) or no memory of seeing the item (N(ovel)). Resting-state fMRI series were acquired three times during the experiment: before encoding (baseline/pre-encoding), immediately following encoding (“post-encoding”) and immediately preceding in-scanner retrieval (“delayed post-encoding”). Functional localizer sessions were run at the end of the experiment. When possible, participants completed the entire experimental protocol twice. The two visits differed only in time of encoding session (morning/evening) and consequently time of in-scanner retrieval which occurred 12h post-encoding. 83 participants completed two full visits.

**Table 1.** Sample characteristics

|  | Young adults |  |  | Older adults |  |  |
| --- | --- | --- | --- | --- | --- | --- |
|  | *n* | Mean (SD) | Range | *n* | Mean (SD) | Range |
| Age | 47 | 26.5 (4.20) | 20-38 | 42 | 67.1(5.81) | 60-81 |
| Education | 45 | 15.2(2.02) | 13-18 | 39 | 16.3(2.06) | 10-21 |
| MMSE | 46 | 29 (1.07) | 27-30 | 42 | 29.1 (1.07) | 27-30 |
| IQ | 46 | 110 (8.97) | 90-125 | 41 | 121 (9.84) | 90-146 |

## Results

### Increase in rsFC between hippocampus and targeted stimulus-sensitive ROIs after encoding

To replicate previous reports of increased rsFC between hippocampus and task-related regions following extensive encoding tasks, we correlated BOLD time series extracted from hippocampus with those from stimulus-sensitive regions, individually defined from the functional localizer protocol (Figure 2A). We then subtracted pre-encoding baseline correlations from post-encoding correlations. The resulting rsFC change measures, one per ROI-pair per participant, were assessed with a linear mixed model fitted as a function of ROI-pair and age group. From the model conducted for hippocampus and the task-related target ROIs; fusiform face area (FFA; sensitive to face stimuli), lateral occipital complex (LOC; objects) and parahippocampal place area (PPA; places/scenes), we observed a post-encoding increase in rsFC compared to baseline for all ROI-pairs (false discovery rate corrected *p*_FDR_ < 0.002) with estimates (change in z-transformed *r*) ranging from 0.08 to 0.12 (Supplementary Table 1, Figure 2B). Excluding 9 observations from 7 participants with LOC <5 vertices did not change the results.

**Figure 2.**
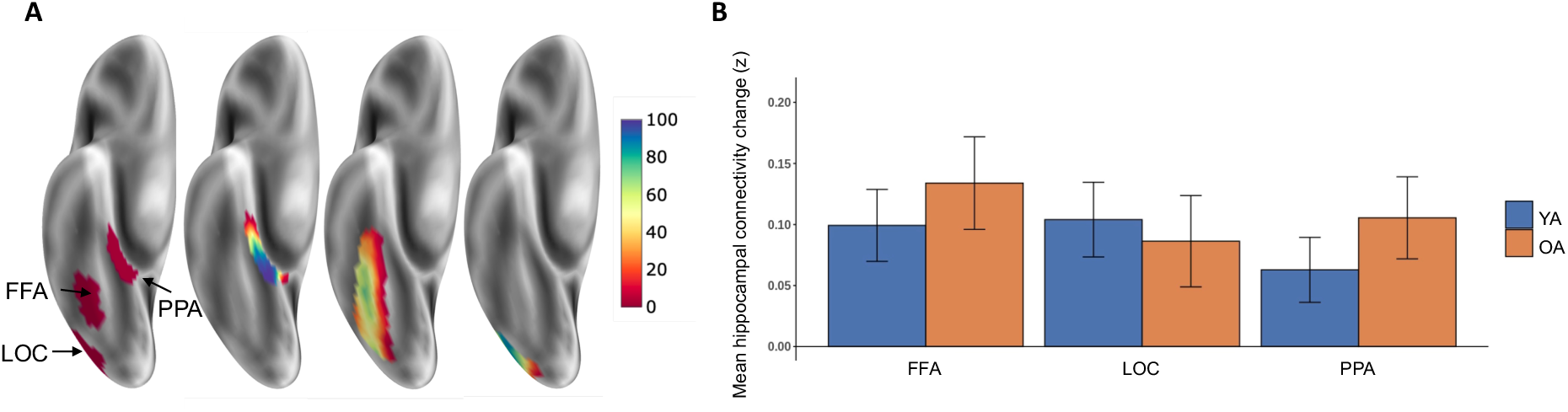
A) Example of category-sensitive regions, defined from the functional localizer protocol (inferior view of the right inflated hemisphere). Right part shows % of participants with a ROI at a given surface vertex. B) Estimated post-pre change in hippocampal rsFC with the three ROIs, separated over age groups (YA/OA = Younger/Older adults). Error bars represent standard errors. FFA: fusiform face area; PPA: parahippocampal place are; LOC: lateral occipital complex.

### Widespread increase in hippocampal-neocortical rsFC after encoding

Next, to assess the extent of post-encoding changes in hippocampal-neocortical rsFC beyond stimulus-category sensitive cortex, we first ran a whole-brain analysis testing for post-pre modulation between the hippocampus and each of 400 nodes in a pre-established neocortical parcellation (24). Following FDR-correction, 305/400 nodes showed significant (*p*_FDR_ < 0.05) post-encoding increases in their rsFC with hippocampus (Figure 3A), indicating that the observed increased coupling with stimulus-sensitive regions occurs as part of an extensive post-encoding modulation of hippocampal functional connectivity, affecting large portions of the cerebral cortex. To test whether the magnitude of this nearly global increase in post-encoding rsFC was unique to the hippocampus, we repeated the analysis after subtracting post-pre rsFC changes between alternative subcortical seed ROIs and the same 400 neocortical nodes (Figure 3B). Most of the observed changes remained significant (*p*_FDR_ < 0.05) after controlling for post-pre rsFC modulations using the following control seed regions: amygdala (201/400 nodes still significant), caudate nucleus (213/400 nodes), putamen (276/400 nodes), and thalamus (237/400 nodes). The uniqueness of hippocampus’ post-encoding behavior was further confirmed by calculating change in the graph-theoretical centrality measure “strength” – the sum of a node’s edges in a weighted graph – for all neocortical and subcortical nodes: here hippocampus showed a numerically larger increase than any other node (Figure 3C). Direct comparisons confirmed that hippocampus’ change in strength was significantly larger than that observed for 391 out of the 404 remaining nodes (including 4 subcortical) in the graph (*p*_FDR_ < 0.05). That is, using rsFC strength as a proxy for centrality in the full brain network, hippocampus showed higher post-pre centrality increase than practically any other node in the brain.

**Figure 3.**
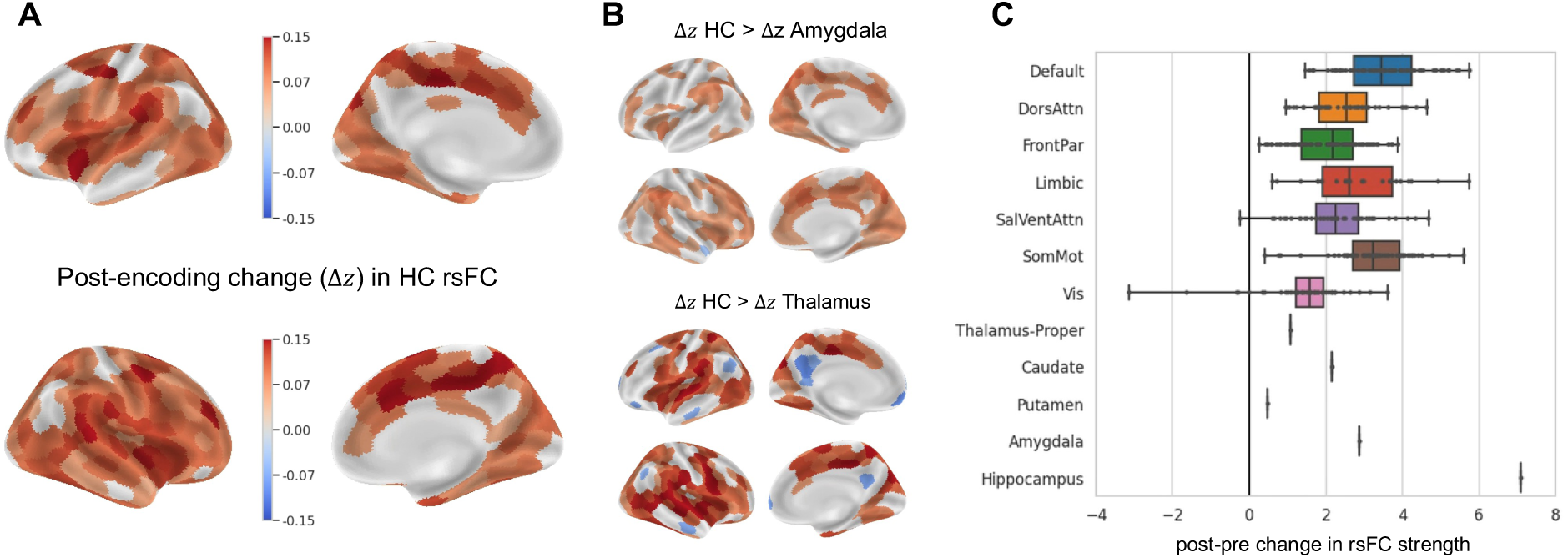
A) 305 nodes showing significant FDR-corrected post-encoding change in rsFC with hippocampus (values represent difference in Fisher-transformed *r*). B) Nodes for which hippocampal rsFC change was significantly (FDR-corrected) different when controlling for change observed using alternative subcortical seed ROIs (Amygdala and Thalamus as examples; see Supplementary Figure 1 for other control seeds). 3) Post-pre rsFC centrality (strength) change per node. Each dot represents a node, neocortical nodes have been arranged into constituent resting-state networks. Boxplot whiskers represent min/max observed nodal value within a network while box limits reflect quartiles; black vertical line represents median strength. SalVentAttn = Saliency/Ventral attention network; DorsAttn = Dorsal attention network; Vis = Visual network; FrontPar = Frontoparietal network; SomMot = Somatomotor network; Default = Default Mode Network.

The nearly global changes in hippocampal rsFC strength following our encoding task were also observed at the network level. A linear mixed model assessing change in rsFC between hippocampus and 7 cortical networks (25) showed rsFC increases with all networks (all *p*_FDR_ < 0.01; Supplementary Table 2, Figure 4A). Highest estimates of rsFC change were seen for the ventral and dorsal attention network while the lowest estimate was associated with the default mode network. Among the subcortical control regions only amygdala showed significant post-encoding rsFC changes at the network level, with dorsal and ventral attention networks and the somatomotor network (*p*_FDR_ < 0.02; Figure 4B).

**Figure 4.**
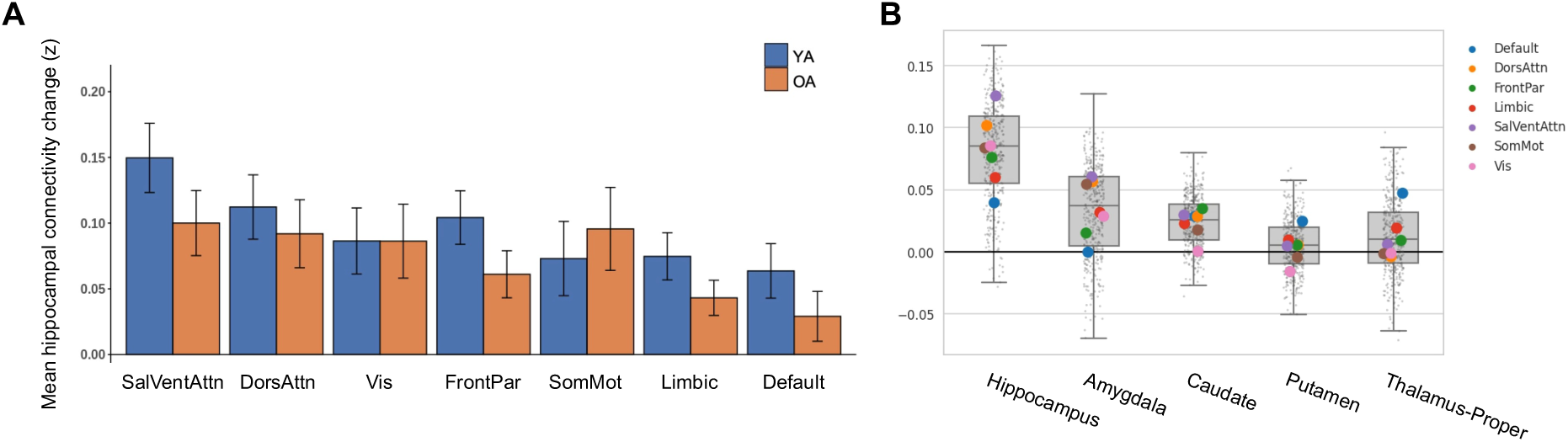
A) Estimated hippocampal rsFC change with 7 neocortical networks, separated over age groups. Errorbars represent standard errors. B) Subcortial seed ROIs change in rsFC with 400 neocortical nodes (gray dots) and networks (colored circles). Box plot whiskers represent ±1.5 interquartile range while box limits reflect quartiles; gray horizontal lines show median changes in *z*.

### Post-encoding change in hippocampal rsFC not present after 12 hours

We next tested the duration of the observed changes in post-encoding hippocampal-neocortical rsFC. We calculated a second rsFC change measure, this time subtracting pre-encoding baseline correlations from hippocampal-neocortical correlations established using resting-state data collected ∼12 hours post-encoding. As participants were scanned over two visits, with baseline resting-state scans once in the morning and once in the evening (and similar for the 12h delayed resting-state scans; see Figure 1), we avoided potential time-of-day effects by averaging change measures from the two visits. Following the same approach as for the original rsFC change analysis reported above, we observed no significant change effects in hippocampal rsFC; not at the single parcel level, nor at the network or stimulus-sensitive ROI level. Moreover, the observed hippocampal rsFC change from baseline to immediate post-encoding rest was significantly greater than the (non-significant) change observed over 12 hours (paired t-test of values extracted from the “global” hippocampal rsFC change mask: *T*(168,5) = 2.48, *p* = 0.014; 192/400 edges with *p*_FDR_ < 0.05 when tested independently). The extensive post-encoding hippocampal rsFC modulations thus appear to be transient in nature, increasing immediately following an intensive learning experience but returning to baseline levels within a 12h timeframe.

### Increases in hippocampal-neocortical rsFC not explained by global signal

To ensure that the nearly global upregulation of hippocampal functional connectivity post-encoding was not driven by residual noise in our data, we compared all subcortical seeds’ baseline correlation against the global gray matter signal. The globally averaged signal (GS) is often considered a measure of spatially diffuse hemodynamic fluctuations of partly non-neuronal origin, understood as noise in the current context (26). If hippocampal dynamics mimic these non-neuronal contributors disproportionally, and the noise-contribution to the global signal increases following encoding, this could theoretically result in apparent increased coupling between HC and gray matter globally. Several of the other subcortical seeds, however, showed significantly stronger baseline correlations with the GS than hippocampus, including both the thalamus (paired t-test; *T*(88) = 7.32; *p* < .001) and caudate nucleus (*T*(88) = 7.00 ; *p* < .001). As none of these nodes showed similar post-encoding rsFC changes as the hippocampus, associations with the global signal cannot explain the current results.

### Post-encoding modulation of hippocampal rsFC mimics encoding patterns

To further characterize the hippocampal-neocortical rsFC change patterns, we compared the spatial profile of modulation observed during post-encoding rest with profiles estimated from encoding and retrieval task periods. Using generalized psychophysiological interaction (gPPI) analysis (27, 28) we established spatial maps of average hippocampal modulation during the two task states (Figure 5A). The specific contrasts used reflected changes in hippocampal functional connectivity during successful source memory operations, controlling for intrinsic connectivity between nodes, and correlated stimulus-induced activation effects. Spatial Spearman correlations between the post-encoding resting-state pattern and gPPI effects observed during active encoding revealed significant similarities in hippocampal-neocortical modulations across the states (*rho* = .427; *p* < .001; Figure 5B). A positive relationship was also found with the retrieval-state gPPI pattern (*rho* = .207; *p* < .001), albeit significantly weaker than the spatial correlation observed with encoding data (*z* = 3.47; *p* = .0005). Additional analyses comparing the observed relationships with permuted null-distributions preserving the spatial autocorrelation structure of the empirical brain maps confirmed significant similarities between encoding-state and post-encoding rsFC modulations (left hemisphere *rho* = .36, *p* = .014; right hemisphere *rho* = .54, *p* < .001; Figure 5C). The retrieval-state gPPI pattern did however not share significant similarities with the post-encoding rsFC change map when controlling for spatial autocorrelations in the data (left hemisphere *rho* = .15, *p* = .31; right hemisphere *rho* = .23, *p* = .10). Thus, global post-encoding changes in hippocampus’ functional connectivity profile preferentially resemble the effect of active encoding.

**Figure 5.**
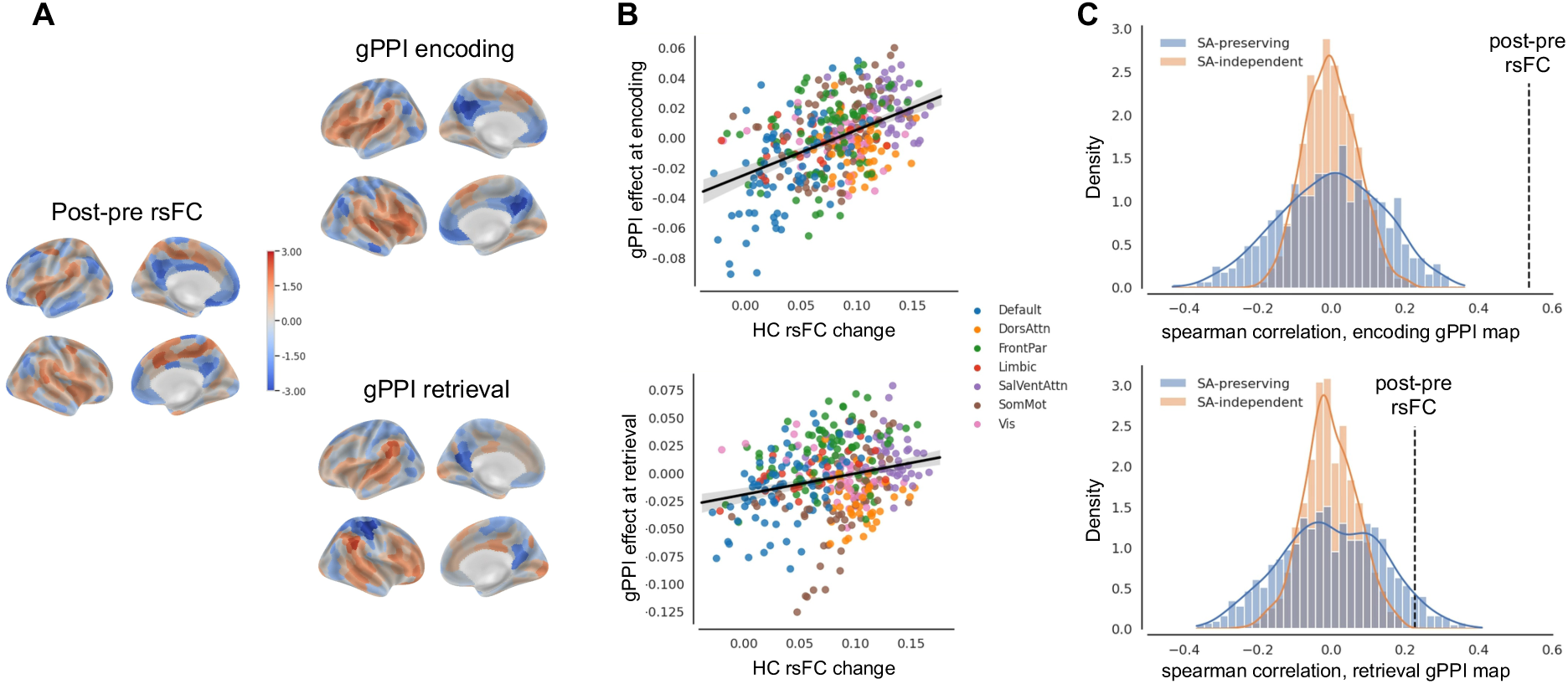
A) Z-standardized maps of average hippocampal functional connectivity changes during resting- and task states. B) Scatterplots showing relationships between task-state encoding (top) and retrieval (bottom) modulations and post-pre rsFC change in hippocampal (HC) functional connectivity with 400 neocortical nodes. Network membership of a node is indicated by its color. C) Permuted null-distributions of spearman correlations between brain maps either ignoring (SA-independent) or incorporating (SA-preserving) spatial autocorrelation (SA) structures in the data. Dashed vertical lines shows empirical correlation between the patterns. Results shown are from right-hemispheric data, similar results were observed in the left hemisphere.

### Post-encoding change in hippocampal rsFC is independent of relevant individual factors

Having established that post-encoding change in HC rsFC involves edges throughout the entire neocortex – in a pattern resembling memory encoding behavior – we tested whether this nearly global modulation was influenced by relevant individual factors. Our main factor of interest, participant age, did however not explain significant variance in the observed reactivation-like connectivity profiles at any resolution or analysis level (category-selective ROI-level: see Supplementary Table 1; network-level: see Supplementary Table 2; neocortical parcel-level: all *p*_FDR_ > .05). Similarly, we estimated average rsFC change within a mask consisting of the 305 nodes showing increased post-encoding rsFC with the hippocampus, i.e. one value per participant. Comparing this “global” hippocampal rsFC change measure between younger and older adults also did not reflect any age differences (Welch separate variances t-test; *T*(86.99) = 0.76, *p* = .45). Unsurprisingly, given their strong correlations with participant age, individual differences in episodic memory performance were also not significantly associated with our measures of hippocampal post-encoding modulation. This was true over a range of tests spanning retention intervals of hours to several days and involving both cued retrieval and forced-choice tasks (Supplementary Table 3).

### Post-encoding Default Mode Network modulations via thalamic interactions

The focus of the current study concerned reactivation-like modulations in hippocampal rsFC. However, considering recent reports of temporal cooccurrence of neurophysiological measures of hippocampal replay and activity increases in the default mode network (DMN) (13, 29), an observation from our control analyses warranted a post-hoc investigation. In our data, hippocampus showed reliable coupling to the DMN both during pre- and post-encoding resting-state periods, i.e., when the periods were investigated in isolation (Supplementary Figure 2). Yet, hippocampal post-pre rsFC change towards the DMN was the weakest observed at the network level (Figure 4A). This was also reflected at the whole-brain parcel level. Here, hippocampus showed increased post-encoding coupling nearly globally across edges, with the conspicuous exception of a set of central DMN regions (medial prefrontal cortex, precuneus, angular gyrus; see Figure 3A). Thalamus, one of the subcortical control seeds, did however show a clear post-encoding modulation of its rsFC with core DMN regions (Figure 6). Moreover, for several DMN nodes this change was significantly stronger than that observed for the hippocampus (see Figure 3B). In line with recent accounts pointing to thalamus as a likely link between hippocampus and the DMN during early consolidation processes (14, 30), this final observation in our data also suggests that parts of the neocortex interact with subcortex – and potentially the hippocampus – via thalamic pathways at post-encoding awake rest.

**Figure 6.**
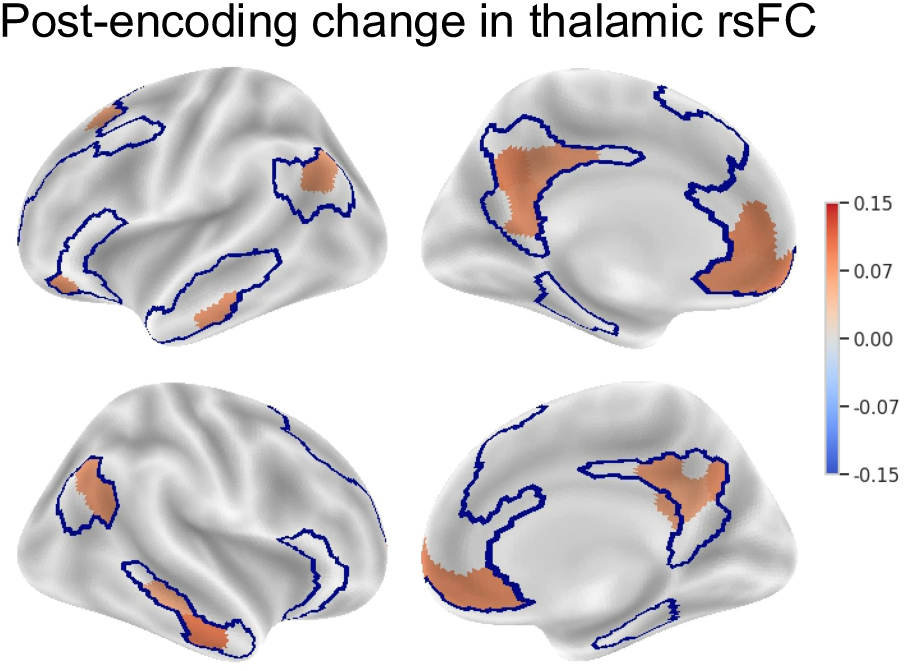
Nodes showing significant FDR-corrected post-encoding change in rsFC with the thalamus (values represent differences in Fisher-transformed *r*). DMN, derived from Yeo’s 7-network parcellation (25), outlined in blue.

## Discussion

We here demonstrate continuation of a memory encoding state into task-free post-encoding rest. Following a period of intensive encoding, hippocampus increased its functional connectivity with large parts of the neocortex – over 75% following corrections for multiple comparisons at the current parcellation resolution. Although these modulations in hippocampal functional connectivity were measured during awake passive rest, their spatial profile across the neocortex resembled hippocampal connectivity patterns observed during active encoding of multimodal stimuli. Importantly, the observed modulations involved regions outside of sensory/perceptual and stimulus-category sensitive cortex; in fact, all the brain’s major functional networks showed some degree of change in their post-encoding hippocampal interactions when contrasted with a pre-encoding baseline.

Such imaging-derived encoding-state continuation into task-free periods has been interpreted as evidence for “reactivation” (7) – potentially reflective of hippocampal-neocortical co-activation patterns seen during neural “replay” in awake resting animals (12, 29), and recently in humans (13). In line with the long tradition connecting hippocampal replay during sharp-wave ripples with memory consolidation processes (31, 32), post-encoding reactivation observed in human fMRI data has reliably been linked with non-declarative (9, 33, 34), and declarative memory processes; the latter through associations with episodic memory performance or detections of stimulus-specific activation patterns (8, 10, 11, 35–39). With few exceptions, however (40), previous approaches investigating reactivation in human participants have been limited to interactions within the medial temporal lobe (MTL) or between hippocampus / MTL-structures and a few selected regions-of-interest.

We here replicate observations of increased post-encoding hippocampal connectivity with face-, place- and object-sensitive regions (e.g., (8)), but also show that these modulations must be understood as part of a nearly global pattern of hippocampal coupling change.

Given the structural similarities between this extensive hippocampal post-encoding coupling change pattern and the global hippocampal-neocortical pattern extracted from the encoding state, we suggest that reactivation may not be limited to reinstatement of relevant sensory characteristics of the encoding task, but also incorporate encoding-relevant processes typically associated with higher-order functional networks, such as attentional allocation, schema integration, and cognitive control (17, 41). In line with this view, previous ROI-focused fMRI-studies in humans have reported post-encoding modulations consistent with integration of novel information in resting-state interactions between hippocampus and ventromedial prefrontal cortical regions (40, 42, 43). Moreover, studies in rhesus monkeys combining electrophysiological recordings and fMRI have found activity increases coinciding with hippocampal ripple events throughout higher-order cerebral cortex (12, 29). Similarly, a recent MEG-study in humans found activity increases source-localized to the parietal lobe (“parietal alpha network”) and DMN regions during hippocampal ripple-events detected during awake rest (13). It should be noted, however, that such ripple-synchronized activity does not indicate causal or direct relationships/connectivity between the hippocampus and neocortical regions. For example, theoretical accounts (30) and fMRI investigations (14) of post-encoding recruitment of the DMN report evidence for thalamic mediation of consolidation-related activity patterns. Our results provide further support for these accounts by the observation of robust functional post-encoding DMN-modulation through thalamic interactions but less so with the hippocampus as seed region.

While investigations of hippocampal interactions with other subcortical structures – e.g. the thalamus – appear promising in understanding the complex dynamics behind brain states supportive of ripple generation and memory reactivation (e.g., (44)), we found it noteworthy that the hippocampus showed the largest post-encoding change in its functional connectivity towards all other nodes in our parcellation, i.e., larger than any other subcortical or neocortical node. The fact that this selective upregulation of hippocampal coupling was absent when estimated from a second resting-state dataset collected ∼12 hours post-encoding supports the current view of hippocampus’ unique role in the initial stabilization of memory traces (e.g., (3)), and resonates with animal studies showing high prevalence of hippocampal replay immediately after an experience, followed by a gradual decay in replay occurrences (4). Our reported measure of hippocampal functional connectivity change is however not a direct reflection of replay or replay-related processes as described in the neurophysiological literature. Although neuronal reactivation in principle can be detected at the BOLD response level (see (7) for a thorough discussion), without simultaneous neurophysiological measurements we cannot know what drives the observed increases in hippocampal coupling. We thus believe the transient centrality boost seen for hippocampus in the post-encoding functional brain network must be interpreted within the context of human imaging-based approaches to reactivation and consolidation processes. Here, post-encoding modulations of pairwise regional BOLD synchronicity have repeatedly been used to predict memory-relevant behavior and through this established a plausible mapping between the methodological approach and the underlying phenomenon of interest, memory consolidation (reviewed in (7)). As we did not observe any associations with memory performance, we cannot draw a similar link between post-encoding functional connectivity and memory-relevant offline processing in our sample. In contrast to previous approaches, however, we here deliberately avoided a limited regions-of-interest analysis approach and consequently had to run strict corrections on all tests of phenotype associations. Moreover, we can still point to several indicators supporting an interpretation of our post-encoding findings in line with early systems consolidation. Most prominent is the similarity with hippocampus-centered encoding patterns, considered the “hallmark” of reactivation (3, 7, 45). Also, the disproportionate increase in hippocampal centrality, relative to all other investigated subcortical and neocortical nodes, indicates strong post-encoding relevance of this core structure for episodic memory formation. To some extent this hippocampal selectivity also points against a pure Hebbian mechanism – sustained novelty-induced neuronal reverberations (46) – behind the current findings as it is unclear why this process should prioritize hippocampal edges. Finally, our observation of increased post-encoding functional connectivity between thalamus and core DMN nodes fits well with recent models of thalamic enabling of low-interference states during hippocampally orchestrated memory reactivation (13, 14, 44).

Similarly, we did not observe any age effects on hippocampal post-encoding modulations. Although this was contrary to our expectations, it suggests that the upregulation of hippocampal-neocortical connectivity after intensive memory encoding represents a universal phenomenon seen across the adult lifespan. Nevertheless, we believe that future studies should continue pursuing this link; one of the most characteristic changes occurring in human cognition with higher age is the decline of episodic memory function (23), but descriptions of awake systems consolidation in samples other than young adults are almost completely absent in the literature (see (22) for a notable exception).

## Conclusion

Is sum, we show that hippocampus’ coupling with the neocortex is upregulated nearly globally during post-encoding awake rest. The spatial configuration of these modulations resembles hippocampal connectivity seen during active encoding, indicating continuation of the hippocampal encoding state into stimulus-free periods. Such systems-wide encoding reinstatement at rest suggests reactivation of memory traces involves aspects beyond perceptual characteristics of encoded events.

## Materials and Methods

### Participants

See Table 1 and Supplementary Methods (section *Participants)* for inclusion criteria and sample characteristics

### Experimental design

Experimental structure is described in Figure 1. One session of the encoding task consisted of 128 trials of an alternating item-face/item-place associative task. Unique items were presented with one of four faces or one of four places (16 trials per unique face/place in a session). Participants were instructed to visualize an interaction between the item and associate and were informed about all subsequent memory tests. All 128 encoded items were presented during the post-encoding memory tests. In-scanner retrieval additionally involved presentation of 64 novel items. Localizer tasks involved 12s stimulus blocks of four categories: faces, places, novel items, or scrambled stimuli. Two stimulus blocks were followed by 12s fixation. During resting-state scans, participants were instructed to focus on a fixation cross. See also Supplementary Methods (sections *Experimental design + Experimental task and stimuli)*.

### MRI acquisition and preprocessing

MRI data were collected on a 3T Siemens Prisma system equipped with a 32- channel head coil. All fMRI data were collected with parameters: TR=1000ms; TE=30ms; flip angle=63°; voxel size=2.5mm isotropic; matrix=90x90; slices=56 (no gap); multiband=4; phase encoding=AP; ascending interleaved acquisition).

Rest/encoding/retrieval runs produced 730 volumes; localizer runs produced 366 volumes. High resolution T1- and T2-weighted volumes were collected for structural descriptions together with field maps for distortion corrections. Data were preprocessed using FMRIPREP (v1.5.3 (47)), as previously described in (48), and denoised following the ICA-AROMA approach (49). See also Supplementary Methods (sections *MRI acquisition + Preprocessing)*.

### ROI definition and estimation of functional connectivity

Category-selective ROIs were defined from localizer data at the individual level using the GLM-contrasts faces>places (FFA), places>faces (PPA), intact>scrambled items (LOC). Bilateral subcortical ROIs were derived from Freesurfer’s automatic segmentation. Neocortical and network parcellations were based on (24) and (25), respectively. All ROIs/parcels/networks were established in participant space. RsFC change measures were calculated by subtracting post-encoding pairwise Fisher-transformed Pearson’s correlations between denoised BOLD timeseries of all ROI/parcel pairs from their pre-encoding correlations. For a given seed, network-level rsFC measures were constructed by averaging over edges involving the seed and parcels assigned to a given network. Task-specific hippocampal FC during encoding and retrieval was estimated using gPPI (28) and consisted of 400 non-directional/symmetrized (27) PPI measures per task state, representing hippocampal-neocortical modulations during successful source memory operations. See also Supplementary Methods (sections *ROI definitions + Preparations for statistical analyses)*.

### Statistical analyses

Linear mixed models (Figure 2B, Figure 4A) were run with Participant ID as random intercepts to account for multiple sessions and included time of day and mean framewise displacement (FD) as covariates. Whole-brain parcel-wise analyses (Figure 3A+B, Figure 4B, Figure 6) were run using linear regressions with seed-parcel rsFC change as dependent variable and FD as covariate. Here, average rsFC change over two visits was used as dependent measure for 69 participants; morning and evening scans were equally distributed in the remaining 20 participants with one valid visit. Nodal strength centrality differences (Figure 3C) were calculated from the individual subjects’ weighted 405x405 neocortical+subcortical rsFC change graphs. Rank-based correlations between rsFC change and task-derived gPPI-patterns (Figure 5) were compared to permuted null-distributions adjusted for spatial autocorrelations (50). All P-values were FDR-corrected for multiple comparisons. See also Supplementary Methods (section *Statistical analyses)*.

### Data availability

## Acknowledgements

This work was supported by The European Research Council (283634, 725025 to A.M.F. and 313440 to K.B.W.); Norwegian Research Council (to A.M.F. and K.B.W.). We thank Prof. Lars Nyberg for helpful comments on the manuscript.

## Supplementary material

### Supplementary Methods

#### Participants

92 participants were enrolled in the present study: 49 younger (20-38 years; 25 females) and 43 older adults (60-80 years; 25 females). All participants were fluent in Norwegian, right-handed, with normal or corrected vision and no history of severe psychiatric or neurological disorder, traumatic or enhanced brain injury, and no current use of medications known to affect the nervous system. Included participants were required to score ≥ 26 on the Mini Mental State Examination (MMSE; (1)), have normal IQ or above (IQ ≥ 85) measured with the Wechsler Abbreviated Scale of Intelligence (WASI; (2)), and no major depression indicated by the Beck Depression Inventory (BDI; (3)) or the Geriatric Depression Scale (GDS; (4)). See characteristics of the final sample in Table 1. All participants signed an informed consent approved by the Regional Ethical Committee of South Norway. The main recruiting strategies included targeted Social Media advertisement, flyers, and posters at selected places (e.g., senior centers). Participants were compensated for the participation.

#### Experimental design

This study is part of a larger project investigating memory consolidation processes at different time scales and their possible relation to memory decline in aging (https://cordis.europa.eu/project/id/725025). Relevant sessions for this report include a fMRI-paradigm consisting of 12 minutes (min) baseline/pre-encoding resting state fMRI (rsfMRI) before two runs of an associative encoding task (12 min each), immediately followed by 12 min post-encoding rsfMRI. Then, approximately 20 minutes after the end of the last encoding run, a forced-choice memory test was administered outside the scanner. Following a 12-hour interval, participants returned for another post-encoding rsfMRI scan before performing three runs in-scanner retrieval (12 min each). Additionally, four functional localizer runs (6 min each), enabling localization of regions sensitive to stimulus categories presented during the encoding task, and several structural scans were performed. After ended scanning, a second and third forced-choice memory test were administered ∼12 hours and ∼5 days post-encoding, respectively. The participants also underwent a session of neuropsychological testing (appr. 3 hours) and several questionnaires. The described structure is depicted in Figure 1.

83 of the 92 participants completed the fMRI-paradigm two times, resulting in 175 sessions. The two visits were separated at least six days apart, with unique sets of task stimuli at each visit. Fifteen sessions had to be excluded for the following reasons: one session from 11 participants were excluded as they reported rehearsing the encoded associations during the post-encoding rest, one of these (a young male) had only one visit and was excluded entirely. Additionally, one session from two participants were excluded due to interruption between encoding and post-encoding rest. One older male was excluded based on motion (mean framewise displacement > 0.2). One young male participant was excluded due to the suspicion that he fell asleep during post-encoding rest scans. Final sample for further analysis was 89 unique participants and 158 sessions. The visits were characterized by either having the encoding task administered in the morning (8 - 10 AM) (n = 79) or in the evening (8- 10 PM) (n = 79). For the morning encoding session participants were instructed to stay awake, i.e., avoid naps, before in-scanner retrieval 12h later. For the evening encoding session, the 12h delay involved sleep at a hospital hotel associated with the scanner facilities. Order of visits was counterbalanced over participants.

Participants were thoroughly trained on the different task components before the experiment began and were informed that their memory for the encoded associations would be tested. All employees administrating the tasks were carefully trained to give identical task instructions across participants.

#### Experimental tasks and stimuli

Stimulus material (from both visits combined) consisted of a total of 384 real-life images of inanimate everyday items, eight images of faces (four males, four females) and eight images of places (four indoor, four outdoor), as well as 384 auditory stimuli in the form of a prerecorded (female voice) name for each item. All item/auditory stimuli were two-syllable Norwegian words. Place- and face stimuli were luminance-matched. A total of 256 items were presented at encoding; of which a predetermined half constituted the task material for participants’ first visit (item images and corresponding auditory item-names 1–128, face images 1–4, and place images 1–4), and the other half of the stimuli constituted the task material for participants’ second visit (i.e., item images and corresponding auditory item-names 129–256, face images 5–8, and place images 5–8). The remaining 128 item stimuli were introduced as novel items during in-scanner retrieval. Apart from the specific images used, the tasks were identical across visits. Training task stimuli consisted of 16 cartoon images and item-names from the same stimuli categories (i.e., items, faces, and places). Item images were obtained mainly from the Bank of Standardized Stimuli (5), some from StickPNG.com and from Google Advanced Image Search under the license “labelled for reuse with modification.” Face images were obtained from Oslo Face Database (described in (6)). Tasks were designed and run using MATLAB 9.7.0 and Psychtoolbox-3 3.0.16.

One session of the rapid event-related encoding task consisted of 128 trials of an alternating item-face/item-place associative task, 64 of each condition. 128 unique items were presented one at a time on the screen for 5 seconds (s) together with either one of four real-life faces, or one of four real-life places (16 trials per unique face/place). Concurrently with the visually presented stimuli, a female voice named the item three times (e.g., “scarf”, “scarf”, “scarf”). Participants were instructed to visualize an interaction between the item and the face- or place-associate before rating the vividness of the imagined interaction on a scale from one to four during a 2s interval following stimulus offset. Finally, a fixation cross appeared and remained on the screen until the beginning of the next trial. Order of conditions and intertrial interval (ITI; 2-7s) was optimized with optseq2 (http://surfer.nmr.mgh.harvard.edu/optseq/). Item-associate combinations were randomized over participants.

The offline memory test was an 8 alternative forced choice (AFC) test where all the items from the preceding encoding task were presented, one at a time, and the participants had to indicate which of the eight possible associates (four faces + four places) they thought the item was paired with at encoding, all in a self-paced manner. This test was performed immediately after post-encoding rest, and after ∼12 hours and ∼5 days.

At in-scanner retrieval, a trial started with the presentation of a visual item and its spoken name (e.g., “scarf”, “scarf”, “scarf”). After a 5s stimulus presentation period, a 2s interval followed in which participants were asked to indicate their recollection of source information associated with the specific item. The four alternatives were: 1) the item was presented together with a face stimulus at encoding; 2) together with a place stimulus at encoding; 3) they remembered seeing the item at encoding but could not recall the associate; 4) the item was not presented at encoding. One session of the retrieval task consisted of 192 trials of which 64 involved an encoding item presented with a face associate, 64 involved an encoding item presented with a place associate, and 64 involved a novel item (i.e., not presented at encoding). Fixation ITI varied between 2-7s and was optimized with optseq2. Item presentation order was randomized over participants with the criterion that an equal number of face-encoded, place-encoded, and novel items were presented within a scanner run (3 runs in total per session).

The localizer task followed an ABN block design (7) in which two 12s stimulus blocks were followed by a 12s fixation block. A stimulus block consisted of continuous presentation of one out of four stimulus categories: faces from the encoding task; places from the encoding task; novel items; scrambled versions of the novel items. During the face and place blocks, one specific face/place stimulus was held static on the screen and participants responded to miniature random changes to the image. During the item/scrambled items block category stimuli were replaced every 1s. Presentation order of stimulus blocks was optimized via custom routines that ensured 1) the same stimulus category was never presented two blocks in a row; 2) over the full session, a specific face/place stimulus were preceded by all place/face stimuli 3) temporal distance between stimulus categories were held as short as possible. Each specific face/place stimuli, as well as the item/scrambled item categories, were presented 8 times over the four runs constituting a localizer session.

During resting-state recordings, participants were instructed to remain awake, keep eyes open and focus on a fixation cross. Afterwards, participants completed a questionnaire of what they were thinking about during scanning.

#### MRI acquisition

Imaging data was collected with a 3T Siemens Prisma MRI unit equipped with a 32- channel Siemens head coil (Siemens Medical Solutions Germany) at Rikshospitalet, Oslo University Hospital, Norway.

Scanning parameters were equal across all fMRI experiments. 56 transversely oriented slices were measured with a BOLD-sensitive T2*-weighted Echo Planar Imaging (EPI) sequence (TR = 1000 ms; TE = 30 ms; flip angle = 63°; matrix = 90 x 90; voxel size = 2.5 × 2.5 × 2.5 mm^3^; FOV = 225 × 225 mm^2^; ascending interleaved acquisition; multiband factor = 4, phase encoding direction = AP). Each encoding, retrieval, and resting state run produced 730 volumes while a localizer run produced 366 volumes. To allow for signal stabilization and avoid T1 saturation effects, the six first volumes of each fMRI run were discarded from the analyses, in addition to the volumes automatically discarded by the Siemens system. Sufficient T1 attenuation was confirmed following preprocessing.

Additional scans included spin-echo field map sequences with opposing phase-encoding directions acquired for distortion correction of the fMRI data; a T1-weighted MPRAGE sequence consisting of 208 sagittally oriented slices (TR = 2400 ms; TE = 2.22 ms; TI = 1000 ms; flip angle = 8°; matrix = 300 × 320 x 208; voxel size = 0.8 × 0.8 × 0.8 mm^3^; FOV = 240 × 256 mm); a T2-weighted SPACE sequence consisting of 320 sagittally oriented slices (TR = 3200 ms; TE = 5.63 ms; matrix = 320 × 300 x 208; voxel size = 0.8 × 0.8 × 0.8 mm3 ; FOV = 256 mm × 240 mm). Additionally, a clinical T2w Fluid-Attenuated Inversion Recovery (FLAIR) sequence was run and inspected by a clinical radiologist.

For the fMRI-sequences, a NordicNeuroLab (NNL; NordicNeuroLab, Norway) 32-inch LCD monitor was positioned behind the scanner and viewed via a mirror attached to the head coil. Participants produced manual responses using a double, two-button NNL ResponseGrip system. Auditory stimuli were presented to the participants with the OptoActive noise cancelling (ANC) II ™ headphones (Optoacoustics, Israel).

#### Preprocessing

We here followed lab-routines that have been described in full in (8). Briefly, we used the Nipype-based (9) FMRIPREP pipeline (version 1.5.3; (10)), with a custom implementation (https://github.com/markushs/sdcflows/tree/topup_mod) of TOPUP distortion correction (11). Quality control of raw+preprocessed data was performed via inspection of visual reports generated by FMRIPREP and MRIQC (12). fMRI data were denoised prior to statistical analysis. Following, non-aggressive removal of ICA AROMA-classified noise components (13), average white matter- and cerebrospinal fluid (CSF) signal timeseries (eroded masks) were extracted from the AROMA-denoised data. Next, using Nilearn routines (14) data were detrended before temporal filtering and regression of WM and CSF timeseries from the AROMA-denoised data, ensuring orthogonality between filters and confound timeseries (15). Finally, the mean signal was added back to the denoised data. Localizer fMRI data were high-pass filtered at 0.005Hz, encoding and retrieval fMRI data were high-pass filtered at 0.008Hz, and resting-state data were bandpass-filtered between 0.008 and 0.09hz

Spatial smoothing (4 mm FWHM) was performed using Freesurfer routines for analyses performed at the surface level (localizer data). No smoothing was applied to data used for ROI/parcel-level analyses.

#### ROI definitions

Functional task-related ROIs were defined individually for each participant based on all available localizer runs from all visits to enable localization of brain areas thought to be especially relevant to the task stimuli in our task, namely face-sensitive FFA, place-sensitive PPA and item-sensitive LOC. A general linear model (GLM) was set up for each participant: the four stimulus categories (faces, places, items, scrambled items) were modeled as blocks of 12s duration according to their presentation schedule while task responses were modeled as stick events. The five task event descriptors were convolved with a canonical (two-gamma) hemodynamic response function (HRF) and added as regressors to the GLM together with their time and dispersion derivatives. Freesurfer FSFAST routines (https://surfer.nmr.mgh.harvard.edu/fswiki/FsFast) were used to estimate parameter estimates and their contrasts from surface level data in fsaverage5 space (10242 vertices per hemisphere)

FFA was defined by surface vertices responding stronger to faces than places within the right posterior and mid fusiform gyrus (16), PPA defined as vertices responding stronger to places than faces within the right parahippocampal gyrus (17) and LOC as vertices responding stronger to items than scrambled items within the right lateral lateral occipital cortex (18), all with an uncorrected threshold of p < 0.0001. If the number of remaining vertices was less than five, the threshold was lowered until five or more contiguous significant vertices were observed. Seven participants had fewer than five LOC vertices at the initial p-threshold. Analyses containing LOC were run with and without these.

The hippocampus and subcortical control ROIs were derived from the Freesurfer automatic subcortical segmentation (19). The selected control regions included thalamus (excluding lateral and medial geniculate bodies), caudate nucleus, putamen and amygdala. All ROIs were bilateral. Caudate and putamen were selected based on observations of modulated functional connectivity in aging and during episodic memory operations involving these structures (8, 20). Amygdala was included as the structure lies adjacent to hippocampus and shares similar MRI signal-to-noise properties (21). Thalamus was included due to suggestions of it having a complementary role to hippocampus during consolidation states (22, 23). All whole-brain parcel-level analyses were based on the Schaefer-Yeo 400 node cortical parcellation (24). Cortical network analyses were based on the Yeo 7- network resting-state parcellation (25). All neocortical parcels/networks were established in participant space through intersections with the Freesurfer-derived cortical ribbon.

#### Preparations for statistical analyses

For preparation of data, statistical analyses, and visualization we used Python 3.7.4, including the use of the packages Scikit-learn (version 0.23.2; (26)), Nilearn (version 0.7.1; (14)), and Pingouin (version 0.3.12; (27)). Linear mixed models were run in R 4.0.0 (R Core Team, 2020) via packages Lme4 (28) and Lmer4Test (29).

The following approach was used to extract rsfMRI BOLD timeseries and estimate functional connectivity measures for a given participant: 1) For the functionally defined ROIs (FFA, PPA, LOC), we ran principal component analysis (PCA) on vertex timeseries from a given ROI to account for the differences in ROI sizes across participants. The timeseries derived from the first PCA component was used as a representative measure of ongoing resting-state fluctuations within the ROI. 2) For the anatomically defined subcortical ROIs, mean timeseries over functional voxels overlapping >50% with the high-resolution structural definition were extracted and averaged over hemispheres. 3) For the whole-brain neocortical 400-node parcellation, functional voxels overlapping >50% with a given neocortical parcel, defined in native high-resolution structural space, and constrained by the cortical ribbon, were considered functional representatives for that parcel and the associated timeseries were averaged. 4) measures of functional connectivity between timeseries were estimated by pairwise Pearson’s correlation for each rest scan separately. The correlation coefficients were Fisher transformed to z-values and post-encoding modulation values were established by subtracting pre-encoding from post-encoding values. 5) Network level measures of functional connectivity change were established by averaging modulation values over all hippocampal edges (or other subcortical seed ROI in the control analyses) with neocortical parcels assigned to a given network (parcel-network correspondence obtained from https://github.com/ThomasYeoLab/CBIG/tree/master/stable_projects/brain_parcellation/Schaefer2018_LocalGlobal).

Task-specific functional connectivity during in-scanner encoding and retrieval was established using generalized psychophysiological interactions (30). For the encoding data, three “psychological” timeseries were set up as boxcar functions, reflecting encoding events (5s duration) characterized by “successful source memory encoding” or “unsuccessful source memory encoding”, as well as response events (2s duration). Memory status was derived from the offline 8AFC test occurring immediately following the post-encoding resting-state scan. For the retrieval data, four psychological timeseries were set up in a similar fashion reflecting “successful source memory retrieval”, “misses” (old items not recognized with correct source information), novel items and response events. Here, memory status was based on the in-scanner responses. Next, denoised task-state BOLD timeseries from hippocampus and 400 neocortical parcels were deconvolved into neuronal estimates (31), and point-by-point multiplied with the demeaned psychological event timeseries (32). The resulting “psychophysiological” timeseries, one per event type, were returned to the BOLD level through convolution with a canonical two-gamma HRF and included in GLMs together with HRF-convolved versions of the psychological functions and the original seed timeseries. As the current investigation focused on hippocampal-neocortical interactions, a total of 800 PPI GLMs was set up for each participant and task-state. 400 of these used a neocortical node as source of the physiological signal in the design matrix and the hippocampal BOLD timeseries as dependent variable. The other 400 used hippocampus as physiological source in the design while the dependent variable spanned all neocortical nodes. Each neocortical node was thus represented on the dependent side in one model and the independent side in another. For our measure of non-directional memory-relevant functional connectivity between hippocampus and a given neocortical node we used the average of the parameter estimates associated with the PPI-regressor representing successful memory trials in the two models including that node. 400 such symmetrized PPI measures (33) representing the full hippocampal-neocortical modulation during source memory operations, were established for both encoding and retrieval state data.

As the experimental paradigm consisted of separate independent memory tests of the encoded content – performed at different delay intervals – we established measures of task-relevant memory performance over four operationalizations: 1) immediate memory: proportion correct item-source assignments on the 8AFC test following immediately after the encoding scan; 2) intermediate memory: proportion correct on the second AFC test ∼12h post-encoding, not including items with wrong source assignment at the immediate test: 3) durable memory: proportion correct on the third AFC test ∼5 days post-encoding, not including items with wrong source assignments in any of the earlier tests; 4) category-level memory; proportion correct face/place assignments out of total exposures to encoded items during in-scanner retrieval, corrected for wrong assignments (e.g., “face” response to item shown with place-associate).

#### Statistical analyses

Linear mixed models were used to test for post-encoding changes in rsFC between hippocampus and the localizer-derived category-sensitive ROIs (Figure 2B), and between hippocampus and the seven cortical networks (Figure 4A). Here, post-pre difference in rsFC was fitted as a function of hippocampal ROI pair (alternatively network pair) and age group. As most participants went through the full paradigm twice, participant ID was added as random intercepts to account for multiple sessions. No global intercept was included in the model, allowing us to assess individual change for each ROI-pair. Estimates of participant motion (mean framewise displacement (FD) over resting-state runs) and time of day (morning or evening scan) were added as covariates. FD, time of day and age group were demeaned. P-values were adjusted for multiple comparisons using False Discovery Rate (FDR)-correction. The same modeling approach was used in control analyses with alternative subcortical ROIs (e.g. Figure 4B).

For the whole-brain parcel-wise analyses (Figure 3A) we ran 400 linear regressions with hippocampal-parcel rsFC change as dependent variable and demeaned FD as covariate. Average functional connectivity change over two visits was used as dependent measure for participants who had been through both morning and evening scans (N=69). In the remaining participants, i.e., those who had only one valid visit (see section *Experimental Design)*, morning and evening scans were equally distributed (N=20; 10 morning scans, 10 evening scans). P-values derived from T-values of the intercepts were FDR-corrected for multiple comparisons and used to assess significance of rsFC change over hippocampal edges. The same approach was used in control analyses with alternative subcortical seed regions, and in the parcel-wise analyses contrasting change in hippocampal-neocortical rsFC with similar measures derived from other subcortical ROIs (Figure 3B). Observing no significant effects of individual motion on the rsFC change measures, seed-parcel rsFC change differences between age groups were tested using independent samples Welch separate variances T-test.

Nodal strength centrality differences were calculated from the individual subjects’ weighted 405x405 neocortical+subcortical rsFC change graphs. A given node’s strength was calculated as the sum of the weights of all edges connected to that node (34). For visualization (Figure 3C) the strength values were averaged across participants and sorted into constituent canonical resting-state networks. Paired T-tests were used to assess differences between hippocampal post-pre strength change and strength changes observed in all other nodes (404 tests in total). Significance was established after FDR-adjustment of p-values.

The comparisons of hippocampal-neocortical post-encoding rsFC change patterns with task-derived gPPI patterns of hippocampal functional connectivity during source memory encoding and retrieval operations (Figure 5) were performed using rank-based (spearman) spatial correlations. Empirical correlations were compared to null-distributions of 1000 spearman correlations where the rsFC change patterns had been randomly permuted. Here, BrainSMASH (35) was used to retain similar spatial autocorrelation structure in the permuted data as in the original, empirical data. Spatial autocorrelation structure was estimated at the surface level, for each hemisphere separately, after calculating geodesic distance between the 200 parcels in a hemisphere using the Python package *surfdist* (*36*).

Global signal timeseries, to be used in control analyses, were extracted from all participants’ baseline rsfMRI scans using a Freesurfer-derived gray matter mask and AROMA-denoised data. For each participant, Spearman correlations between the global signal and BOLD-timeseries from subcortical ROIs during the same baseline scan were estimated and Fisher-transformed before being averaged across visits. Group-level differences between subcortical structures’ baseline correlation with the global signal were tested using paired-samples t-tests.

Associations between individual differences in memory performance and post-encoding hippocampal rsFC modulations were tested over four operationalization of retention success, using linear mixed models at the category-selective ROI level and network level, and partial Spearman correlations at the whole-brain parcel level. A linear mixed models was run for each hippocampal ROI pair (alternatively network pair) separately; here, memory performance was fitted as a function of rsFC change and age group, participant ID included as random intercepts to account for multiple visits, and time of day added as covariate. The partial correlations were run iteratively over hippocampal rsFC change with the 400 neocortical parcels, including participant age as covariate. We also estimated rsFC change within a mask consisting of the 305 nodes showing increased post-encoding rsFC with the hippocampus (i.e., one value per participant), and compared this “global” hippocampal-neocortical change measure with our memory performance measures using a similar partial correlation approach. All P-values were FDR-corrected for multiple comparisons. As a few participants missed data on some of the memory tests, the number of observations entered in the analysis varied slightly between operationalizations. Sample sizes for the different tests are reported in Supplementary Table 3.

---

**Supplementary Figure 1.**
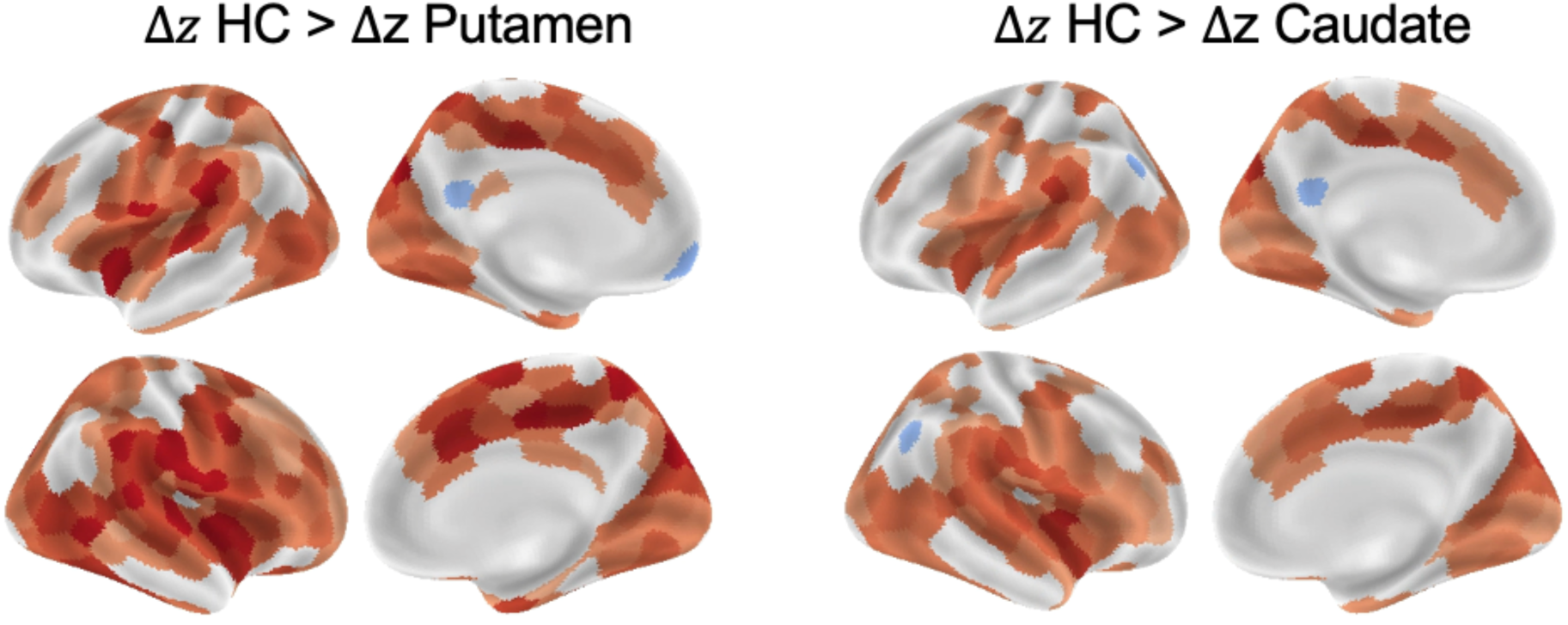
Nodes for which hippocampal post-pre rsFC change was significantly (FDR-corrected) different when compared to change observed using alternative subcortical seeds.

**Supplementary Figure 2.**
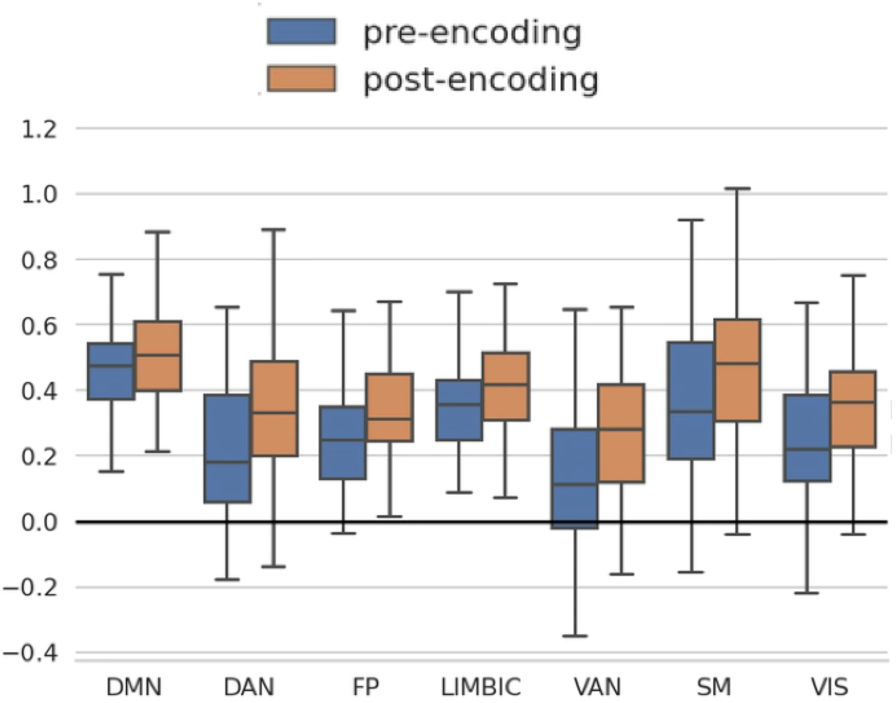
Mean hippocampal rsFC over neocortical networks, estimated separately for pre-encoding and the post-encoding scans. DAN = Dorsal Attention Network; FP = Frontoparietal Network; VAN = Ventral Attention Network; SM = Somatomotoric Network; VIS = Visual Perceptual Network. Boxplot whiskers represent the 1.5 interquartile range.

**Supplementary Table 1.** Linear mixed model results from tests of hippocampal (HC) post-encoding rsFC change with category-sensitive ROIs

|  | Difference in rsFC post-encoding vs pre-encoding |  |  |  |
| --- | --- | --- | --- | --- |
| *Predictors* | *Estimates* | *95% CI* | *t-value* | *pFDR* |
| HC-FFA | 0.12 | 0.07 – 0.17 | 4.57 | **<0.001** |
| HC-LOC | 0.10 | 0.05 – 0.15 | 3.78 | **0.001** |
| HC-PPA | 0.08 | 0.03 – 0.13 | 3.29 | **0.002** |
| Age group (demeaned) | 0.03 | -0.06 – 0.12 | 0.71 | 0.478 |

**Supplementary Table 2.** Linear mixed model results from tests of hippocampal (HC) post-encoding rsFC changes with canonical resting-state networks

|  | Difference in HC rsFC pre-encoding vs post-encoding |  |  |  |
| --- | --- | --- | --- | --- |
| *Predictors* | *Estimates* | *95% CI* | *t-value* | *pFDR* |
| Default | 0.04 | 0.01 – 0.08 | 2.31 | 0.023 |
| Dorsal Attention | 0.10 | 0.06 – 0.14 | 5.19 | **<0.001** |
| Frontoparietal | 0.08 | 0.04 – 0.12 | 4.21 | **0.001** |
| Limbic | 0.06 | 0.02 – 0.09 | 2.96 | **0.004** |
| Somatomotor | 0.08 | 0.04 – 0.12 | 4.21 | **0.001** |
| Ventral Attention | 0.12 | 0.09 – 0.16 | 6.42 | **<0.001** |
| Visual | 0.08 | 0.05 – 0.12 | 4.35 | **<0.001** |
| Age group (demeaned) | 0.01 | -0.05 – 0.07 | 0.27 | 0.790 |

**Supplementary Table 3.** Results of tests of associations between individual differences in memory performance and post-encoding rsFC modulations. Performance-columns (YA/OA = Younger/Older adults) show mean correct source memory with standard deviations in parenthesis. Partial correlation results column (whole-brain + global mask analysis) show results without (outside parenthesis) and with participant age (inside parenthesis) included as covariate.

|  | HC rsFC vs performance |  |  |  |  |  |
| --- | --- | --- | --- | --- | --- | --- |
| *Memory performance operationalization* | *YA performance* | *OA performance* | *Whole-brain (400 edges), Partial spearman* | *Yeo 7 networks, Linear mixed model* | *Localizer ROIs Linear mixed model* | *Global mask Partial spearman* |
| Immediate memory (~1h, 8AFC) | N=46<br>74.0% (16.8) | N=42<br>54.6% (18.7) | top $\rho = .35$ (.32)<br>all $pFDR > .05$ | top estimate = 14.31<br>all $pFDR > .05$ | top estimate = 2.82<br>all $pFDR > .05$ | $\rho = -.02$ (-.12)<br>$p = .85$ (.28) |
| Intermediate memory (12h, 8AFC) | N=46<br>64.1% (21.0) | N=42<br>39.5% (19.4) | top $\rho = .29$ (.26)<br>all $pFDR > .05$ | top estimate = 10.89<br>all $pFDR > .05$ | top estimate = 1.19<br>all $pFDR > .05$ | $\rho = -.03$ (-.11)<br>$p = .82$ (.31) |
| Durable memory (5d, 8AFC) | N=45<br>52.8% (22.3) | N=42<br>28.8% (16.0) | top $\rho = -.32$ (-.30)<br>all $pFDR > .05$ | top estimate = 2.42<br>all $pFDR > .05$ | top estimate = -2.40<br>all $pFDR > .05$ | $\rho = -.06$ (-.17)<br>$p = .58$ (.12) |
| Category-level memory (12h, in-scanner) | N=47<br>70.1% (19.7) | N=42<br>48.8% (23.5) | top $\rho = -.30$ (-.29)<br>all $pFDR > .05$ | top estimate = 15.59<br>all $pFDR > .05$ | top estimate = 5.79<br>all $pFDR > .05$ | $\rho = -.03$ (-.12)<br>$p = .74$ (.25) |

